# Virucidal efficacy of different formulations for hand and surface disinfection targeting SARS CoV-2

**DOI:** 10.1101/2020.11.08.373738

**Authors:** Katrin Steinhauer, Toni Luise Meister, Daniel Todt, Adalbert Krawczyk, Lars Paßvogel, Britta Becker, Dajana Paulmann, Birte Bischoff, Maren Eggers, Stephanie Pfaender, Florian H H Brill, Eike Steinmann

## Abstract

In the ongoing SARS CoV-2 pandemic effective measures are needed, and guidance based on the methodological framework of the European committee for standardization (CEN) can help to choose effective disinfectants on an immediate basis. This study demonstrates that two commercially available formulations for surface disinfection and one formulation for hand disinfection claiming *“virucidal activity against enveloped viruses”* are effectively inactivating SARS-CoV-2. This study emphasizes that chemical disinfectants claiming *“virucidal activity against enveloped viruses”* are an effective choice to target enveloped SARS-CoV-2 as a preventive measure.

## Introduction

On 30 January 2020 WHO declared the outbreak a novel coronavirus designated SARS-CoV-2 a public health emergency of international concern (PHEIC), being WHO’s highest level of alarm [1]. Throughout the ongoing SARS-CoV-2 pandemic from December 2019 through 11 October 2020 over 37 million cases of COVID-19 and 1 million deaths have been reported on a global basis. Recently, within only 1 week (5 October through 11 October 2020) over 2.2 million new cases of SARS-CoV-2 infections and 39.000 deaths associated with COVID-19 were reported, being the highest number of cases so far in the ongoing SARS-CoV-2 pandemic [2].

In order to prevent further spreading of SARS-CoV-2, the WHO recommended hygiene measures such as the use of 70 % ethanol [3]. To enable the use of other suitable disinfectants, the German Robert-Koch-Institute has been recommending the use of disinfectants claiming at least “virucidal activity against enveloped viruses” in the context of the SARS-CoV-2-outbreak [4].

This recommendation is based on the methodological framework of the European Committee for Standardization (CEN): CEN has defined a set of surrogate test organisms, which are representative for certain groups of microorganisms. A proven efficacy against these representative surrogate test organisms allows efficacy claims for the respective group of organisms’ e.g. bactericidal, yeasticidal, fungicidal or virucidal efficacy [5]. For the claim “virucidal activity against enveloped viruses”, vaccinia virus has been defined as the relevant surrogate organism.

As such, disinfectants and antiseptics claiming “virucidal activity against enveloped viruses”, based on the methodological framework of CEN, can be claimed effective against all enveloped viruses including coronaviruses such as SARS-CoV-2 [6]. This holds also true for disinfectants claiming “virucidal activity against enveloped viruses” based on the German DVV/RKI guideline [7].

Despite this guidance throughout the ongoing SARS-CoV-2-pandemic question often arises, of whether a certain formulation has proven efficacy against SARS-CoV-2. Therefore, this study aimed to investigate the efficacy of three different typical formulations used for hand or surface disinfection against SARS-CoV-2 using the European Standard EN 14476 protocol. Efficacy data using SARS-CoV-2 were compared to data obtained with the surrogate test virus vaccinia as defined in EN 14476 and the comparable German DVV/RKI guideline, respectively.

## Material and Methods

### Tests strains and cultivation

Test virus suspensions were prepared by infecting susceptible cells with different multiplicities of infection (MOI). For Modified vaccinia virus Ankara (provided from the Institute of Animal Hygiene and Veterinary Public Health of the University Leipzig), BHK-21 cells were used (provided by Friedrich Löffler institute); for vaccinia virus Elstree (kindly provided by Prof. Sauerbrei, University of Jena, Jena, Germany), CV1 cells (kindly provided by Prof. Sauerbrei, University of Jena, Jena, Germany) were used. SARS-CoV-2 (strain Essen) was propagated on Vero E6 cells as previously described [8].

### Quantitative Suspension tests according to EN 14476 or DVV/RKI guideline

Quantitative suspensions tests were carried out as described in EN 14476 [6] or in the DVV/RKI guideline [7], and the respective test protocol used is indicated for each data set. Briefly, efficacy of three commercially available disinfectants was studied against vaccinia virus (strain modified vaccinia virus Ankara (MVA) ATCC VR −1508 or vaccinia virus, strain Elstree) and SARS-CoV-2 using The virus suspension was added to the product test solution and the interfering substance. A virus control mixture was also assessed using distilled water in place of the test product. After the specified contact time indicated in table 1, virucidal activity of the solution was immediately suppressed by dilution with nine volumes of ice-cold medium (MEM + 2.0% FCS) and serially diluted 10-fold. Due to the immediate titration, no after-effect of the test product could occur. For each test suspension, 6 wells of a microtitre plate containing a confluent monolayer of the respective host cells were inoculated with 100 μL of test suspension, and the cells were incubated at 37°C in a humidified atmosphere under 5% CO_2_.

**Table 1:** Comparison of log_10_ reduction of SARS CoV-2 and vaccinia viral titres by three different formulations used for either surface or hand disinfection. Experiments indicating a ≥ 4 log_10_ reduction of viral titer are given in bold. ^a^Test was carried out according to the DVV/RKI guideline using 10 % fetal calf serum as organic load instead of 0,03% bovine serum albumin. ^b^Test virus used: strain modified virus Ankara (MVA) ATCC VR −1508 ^c^ Test virus used: strain Elstree ^d^Test was carried out according EN 14476 using 0,03% bovine serum albumin as organic load. ^e^RF value was acquired by large volume plating. *pre-screening experiments: data is based on n=1; due to cytotoxicity of the tested substance the detection limit did not allow to detect higher virus reduction. n.d.: not determined

| Formulation | Concentration (% v/v) | Contact time (sec.) | Titre of the virus control (log<sub>10</sub> TCID<sub>50</sub>/ml) SARS-CoV-2 | Logarithmic Reduction Factor (RF) SARS-CoV-2 | Titre of the virus control (log<sub>10</sub> TCID<sub>50</sub>/ml) Vaccinia virus | Logarithmic Reduction Factor (RF) Vaccinia virus |
| --- | --- | --- | --- | --- | --- | --- |
| A | 20 | 15 | 6.22 | **≥4.02** | 7.44<sup>a,b</sup><br>± 0.41 | 0<sup>a,b</sup><br>± 0.40 |
| A | 20 | 30 | n.d. | n.d. | 7.76<sup>a,b</sup><br>± 0.39 | 0.38<sup>a,b</sup><br>± 0,40 |
| A | 80 | 15 | 6.22 | **≥4.02** | n.d. | n.d. |
| A | 90 | 15 | n.d | n.d. | 7.44<sup>a,b</sup><br>± 0.41 | **≥ 4.25<sup>a,b</sup>**<br>± 0.28 |
| A | 90 | 30 | n.d | n.d. | 7.76<sup>a,b</sup><br>± 0.39 | **≥ 4.25<sup>a,b</sup>**<br>± 0.28 |
| B | 10 | 60 | n.d. | n.d. | 7.75<sup>a,c</sup> ±<br>0.33 | 0,25<sup>a,c</sup><br>± 0,48 |
| B | 20 | 15 | 6.22 | **≥4.02** | n.d. | n.d. |
| B | 20 | 30 | 6.37 | ≥ 3.17* | n.d. | n.d. |
| B | 20 | 60 | 6.37 | ≥ 3.17* | n.d. | n.d. |
| B | 80 | 15 | 6.22 | **≥ 4.38<sup>e</sup>** | n.d. | n.d. |
| B | 80 | 30 | 6.37 | **≥ 4.38<sup>e</sup>** | 7.82<sup>a,c</sup><br>± 0.37 | **≥ 4.32<sup>a,c</sup>**<br>± 0,26 |
| B | 80 | 60 | 6.37 | ≥ 2.17* | 7.82<sup>a,c</sup><br>± 0.37 | **≥ 4.51<sup>a,c</sup>**<br>± 0,31 |
| C | 20 | 15 | 6.22 | **≥ 4.02** | 7.67<sup>b,d</sup><br>± 0.33 | 0,17<sup>b,d</sup><br>± 0,58 |
| C | 20 | 30 | 6.22 | ≥ 3.02* | n.d. | n.d. |
| C | 80 | 15 | 6.22 | ≥2.02* | 7.67<sup>b,d</sup><br>± 0.33 | **≥ 4.19<sup>b,d</sup>**<br>± 0.33 |
| C | 80 | 30 | 6.22 | **≥ 4.38<sup>e</sup>** | n.d. | n.d. |

n.d.: not determined

After incubation, the cells were examined microscopically for infectivity and cytopathic effects (CPE). The virus titers were expressed as tissue culture infectious dose 50% (TCID50/mL). The virucidal activity was determined asthe difference betweenmthe logarithmic titer of the virus control minus the logarithmic titer of the test virus (log_10_ TCID50/mL). This difference was given as a RF including its 95% confidence interval. A reduction in virus titer of ≥4 log_10_ (corresponding to an inactivation of ≥99.99%) was regarded as evidence of sufficient virucidal activity. The calculation was performed according to EN14476 [6] or DVV/RKI guideline, respectively [7].

A ready-to-use alcohol-based surface disinfectant designated formulation A (trade name: mikrozid universal; 100 g contains: 17.4 g propan-2-ol, 12.6 g ethanol (94%); Schülke & Mayr GmbH, Germany) was used as one test formulation. In addition, a QAC-based formulation for surface disinfection was used, containing quaternary ammonium compounds designated formulation B (trade name: mikrozid sensitive; 100 g contains: 0.26 g Alkyl(C12-16)dimethylbenzylammonium-chloride (ADBAC/BKC (C12-16)); 0.26 g Didecyldimethylammoniumchloride (DDAC), 0.26g Alkyl(C12-14)ethylbenzyl-ammoniumchloride (ADEBAC (C12-14)). As a third formulation an alcoholic hand disinfectant based on propan-2-ol was used (trade name: desmanol pure designated formulation C (100 g contains: 75 g propan-2-ol). Disinfectant concentrations and contact times used throughout this study were based on the existing “virucidal efficacy against enveloped viruses” efficacy claims for the three disinfectants and are indicated. Experiments were carried out under conditions of low organic soiling (0.3 g/L bovine serum albumin (BSA); “clean conditions”) as defined in EN 14476 [6] or in the presence of 10% fetal calf serum as defined in the DVV/RKI guideline [7].

All experiments were carried out as independent experiments and data presented are based on at least two experiments. Validation controls as defined in the test protocols (EN 14476 or DVV/RKI-guideline) were found to be effective in all experiments indicating validity of presented data.

## Results and Discussion

Throughout the ongoing SARS-CoV-2 pandemic, effective disinfection protocols are needed to support prevention strategies worldwide. Thus, we investigated three different disinfectant formulations in regard to their effectiveness against SARS-CoV-2, including two formulations for surface disinfection and one hand disinfectant. Formulations were based on either alcohol or quaternary ammonium compounds (QACs) with known efficacy against the enveloped vaccinia virus (strain modified vaccinia virus Ankara or strain Elstree, respectively) as established in the European Standard EN 14476 and the quite similar German DVV/RKI guideline. [6,7]. In both test protocols a logarithmic reduction of the test virus by at least 4 log_10_ is required to claim “virucidal activity against enveloped viruses”. Data obtained for SARS CoV-2 by using the EN 14476 test protocol in comparison to data obtained for vaccinia virus using either the DVV/RKI or the similar EN 14476 test protocol are summarized in Table 1. In preliminary screening experiments due to the cytotoxicity of the tested substance the limit of detection did not allow to verify the 4 log_10_ requirement of EN 14476. Thus, further experiments were carried out with either lower concentrations and / or the use of large volume plating to the enlarge the detectability threshold.

Formulation A (alcoholic surface disinfectant) effectively inactivated SARS-CoV-2 by ≥ 4,02 log_10_ within 15 seconds already at a 20% (v/v) dilution. In comparison, formulation A was not found to be effective under these conditions when using the surrogate test virus MVA. Here, a RF ≥ 4.25 log_10_ was obtained, when using the higher test concentration of 90% (v/v), indicating a higher stability of MVA to formulation A compared to SARS-CoV-2.

Formulation B was also found to be effective against SARS-CoV-2 under the chosen test parameters, indicated by ≥ 4 log_10_ RF within 15 seconds at a concentration of 20% and 80% (v/v), respectively. When using the surrogate test virus vaccinia strain Elstree, formulation B was found to be effective within 30 seconds contact time. Interestingly, this formulation was found to be ineffective against vaccinia strain Elstree when tested in a 10 % (v/v) dilution within 60 seconds based on preliminary data from our lab, i.e. RF = 0,25 log_10_. However, for this formulation further data are needed to evaluate, whether this formulation would be effective against vaccinia strain Elstree meeting the 4 log_10_ requirement also within 15 seconds.

Formulation C was found to give ≥ 4,02 log_10_ RF within 15 seconds at 20 % (v/v) when using SARS-CoV-2 as a test virus. For MVA only the 80% (v/v) concentration was found to result in a ≥ 4,19 log_10_ RF, whereas the 20 % (v/v) concentration was not found to equally inactivate MVA, indicated by 0,17 log_10_ RF.

The data presented in this study indicate that the enveloped SARS-CoV-2 is more susceptible to the tested alcoholic biocidal formulations (A and C) compared to the enveloped MVA, which has been established as a surrogate standard test virus in European and German test protocols. For QAC-based formulation B, our data also indicate that SARS-CoV-2 is at least equally susceptible compared to the standard test virus vaccinia strain Elstree. Preliminary data from our lab even indicate a more limited stability of SARS-CoV-2 to the QAC-based formulation when compared to vaccinia virus, which needs to be verified in future studies. This finding is in good agreement with recently published data indicating a good efficacy of QAC-based formulations against three different SARS-CoV-2 strains within 30 s contact time [9].

In conclusion, data from our study undermine the validity of the surrogate test strain concept as established by national and international institutions such as the DVV in Germany and the European Standardization Committee (CEN). This is in good alignment with earlier published data investigating the chemical susceptibility of the human pathogen *Candida auris* compared to the surrogate test organism *Candida albicans* [10]. In the present study as well as in the above mentioned earlier study the surrogate test virus and the surrogate test yeast, respectively were found to be more resistant to the applied chemical disinfectants then the targeted outbreak organism.

Thus, based on the surrogate concept chemical disinfectants claiming “virucidal activity against enveloped viruses” will be an effective choice to target enveloped SARS-CoV-2 as a preventive measure.

